# AID Shapes Proliferation and Cell-of-Origin-associated Transcriptional Programs in Diffuse Large B-cell Lymphoma

**DOI:** 10.64898/2026.02.24.707639

**Authors:** Linda H. Gijsbers, Tijmen P. van Dam, Martin F.M. de Rooij, Guus de Wilde, Richard J. Bende, Marcel Spaargaren, Ad van Gorp, Carel J.M. van Noesel, Jeroen E.J. Guikema

## Abstract

Activation-induced cytidine deaminase (AID), which is essential for antibody diversification, exhibits elevated expression in the activated B-cell-like (ABC) subtype of diffuse large B-cell lymphoma (DLBCL). Here, we demonstrate that AID modulates transcriptional programs linked to cell cycle progression, proliferation, and DLBCL subtype identity. AID loss in ABC-type DLBCL cell lines negatively impacts MYC and E2F pathway activity, while AID re-expression restores activity, establishing a causal link. Consequently, loss of AID delays G1/S cell cycle transition and reduces proliferation. In addition, AID expression skews transcriptional programs towards ABC-type DLBCL in cell lines. In agreement, AID expression correlates with ABC-type gene expression in primary DLBCL patient samples. Moreover, AID overexpression resulted in increased IRF4 protein levels, and enhanced NF-κB activity, supporting AID’s role in reinforcing the ABC-type identity. Shared enrichment of the IRF4 co-factor BATF in AID-high tumors of both ABC- and GCB-subtypes points towards a common mechanism driving subtype skewing. These findings underscore a broader role for AID in DLBCL pathogenesis, establishing AID as a key regulator of transcriptional programs linked to cell cycle progression and DLBCL subtype.

## INTRODUCTION

Activation-induced cytidine deaminase (AID) is a crucial enzyme in B-cell biology, playing a central role in antibody diversification processes. In normal B cells, AID is primarily responsible for initiating somatic hypermutation (SHM) and class switch recombination (CSR) of immunoglobulin genes (1, 2). These processes are essential for generating high-affinity antibodies and switching antibody isotypes respectively, thereby enhancing the adaptive immune response (3).

Beyond its canonical functions, AID has been implicated in processes that extend its influence beyond immunoglobulin genes. One such role is its potential involvement in active DNA demethylation, where AID may deaminate 5-methylcytosine to thymine, initiating a base excision repair process that results in unmethylated cytosine (4–8). This suggests that AID could have a broader effect on gene expression by modulating the epigenetic landscape. AID can therefore influence methylation diversity in genomic regions important during the germinal center (GC) stage of B-cell maturation (8).

In B-cell malignancies, the role of AID is more complex and potentially oncogenic, especially in diffuse large B-cell lymphoma (DLBCL), the most common B-cell non-Hodgkin lymphoma (B-NHL), which shows marked genetic and clinical heterogeneity (9–13). The disease is typically classified into two main molecular subtypes: activated B-cell-like (ABC) and germinal center B-cell-like (GCB), each with distinct gene expression profiles and clinical outcomes (9, 12, 14), where ABC-type DLBCL generally has a worse clinical prognosis than GCB-type DLBCL (9, 11, 14). It has been shown that AID is expressed in both GCB- and ABC-type DLBCL, but its expression levels are significantly higher in the ABC subtype (15–18). This higher expression in ABC-DLBCL is somewhat unexpected, as this subtype is generally characterized by down-regulation of germinal center-specific genes. The elevated AID expression in ABC-DLBCL may be attributed to the constitutive activation of the NF-κB pathway and high expression of IRF4, which are established regulators of AID expression in GC B cells (19–23). Expression of AID has been associated with poor prognosis in DLBCL in several studies, and its activity has been linked to increased genomic instability and the generation of oncogenic translocations (15, 24–28).

Evidence from mouse models underscores the role of AID in driving lymphomagenesis. Transgenic mice with constitutive expression of AID develop B-cell malignancies, indicating that deregulated AID expression can induce tumorigenesis or cause tumor progression (29–31). For instance, in the λ-MYC mouse model, in which MYC expression is driven by immunoglobulin light chain regulatory elements, AID expression is crucial for the development of DLBCL-like disease (32). In addition, AID-deficient mice indicate that AID is required for lymphomagenesis and tumor progression (32–36), as Eµ-c-myc transgenic mice lacking AID expression fail to form GC-derived B-cell lymphomas but instead develop pre-B cell tumors (34). AID-deficiency also prevents the development of GC-derived Bcl6-dependent lymphomas (32). Furthermore, in an IL6-induced tumor model of Burkitt’s lymphoma, it was shown that AID is required for *c-myc/IgH* chromosome translocations, a crucial driver event in disease pathogenesis (33). This evidence highlights the oncogenic potential of AID when its expression is dysregulated, further supporting its role in lymphoma development.

AID has been shown to influence the expression of non-immunoglobulin genes, potentially through its role in DNA demethylation or other epigenetic mechanisms (8, 36). In a recent publication, Gómez-Escolar *et al.* showed that B-cell lymphomas in the λ-MYC mouse model that experienced AID expression showed a different transcriptional profile than non-AID-experienced tumor cells (35). Furthermore, Teater *et al.* showed that overexpression of AID in a Bcl2-driven murine B-cell lymphoma model induces DNA methylation heterogeneity, resulting in a more aggressive disease (36). These findings suggest a potential role for AID in shaping gene expression in malignant B cells through methylome alterations.

Given these observations, we set out to investigate the broader impact of AID on gene expression profiles in human DLBCL. In this study, loss and gain of AID expression in DLBCL cell lines revealed that AID predominantly modulates proliferation-associated transcriptional programs, highlighting a previously underappreciated regulatory role in lymphoma biology. By integrating these *in vitro* data with gene expression profiles from primary DLBCL samples, we show that AID expression levels associate with distinct proliferation-related gene signatures and cell-of-origin-linked programs, suggesting that AID contributes to the molecular heterogeneity of DLBCL. Our findings support a model in which AID not only drives lymphomagenesis through its canonical mutagenic activity but also shapes proliferative and cell-of-origin-associated transcriptional landscapes, with potential implications for risk stratification and therapeutic targeting in this heterogeneous disease.

## MATERIALS AND METHODS

### Cell culture

Cell lines HBL1, TMD8, (kind gifts from Prof. Georg Lenz, University of Muenster, Germany) OCI-LY7, OCI-LY10, OCI-LY18 (kind gifts from Prof. Ulf Klein, University of Leeds, UK), SU-DHL-6 (ATCC, Manassas, VA) were cultured in IMDM (Invitrogen Life Technologies, Carslbad, CA) supplemented with heat-inactivated fetal calf serum (FCS), 2 mM L-glutamine, 100 U/ml penicillin and 100 ug/ml streptomycin (pen/strep) (Gibco, Thermo Fisher Scientific, Waltham, MA). ABC-type DLBCL cell lines TMD8 and HBL1 were maintained in 20% FCS (HyClone, GE Healthcare Life Sciences, Pittsburgh, PA) and GCB-type cell lines OCI-LY18 and SU-DH-L6 in 10% FCS. OCI-LY10 was cultured in presence of 20% human serum (Sigma Aldrich, St. Louis, MO). HEK293T and Phoenix-GaLV cells (ATCC) were cultured in DMEM (Invitrogen Life Technologies) supplemented with 10% FCS, 2 mM L-glutamine, 100 U/ml penicillin, 100 μg/ml streptomycin.

### Plasmid constructs and retroviral/lentiviral transductions

The pL-CRISPR.EFS.GFP plasmid (Addgene #57818), a gift from Benjamin Ebert, was used to generate AID knockout DLBCL clones. The following gRNAs were used for AID targeting: CTTTGGTTATCTTCGCAATA and GGTCCCAGTCCGAGATGTAG. Control clones were established using the same vector lacking a gRNA insert. Lentiviral particles were produced by co-transfecting HEK293T cells with the CRISPR plasmid, psPAX2 packaging plasmid (Addgene #12260), and pMD2.G (VSV-G envelope plasmid; Addgene #12259), gifts from Didier Trono, at a molar ratio of 4:2:1, respectively. Transfections were performed using Genius transfection reagent (Westburg, Leusden, The Netherlands) following the manufacturer’s instructions. Medium was refreshed after one day and viral supernatants were harvested after 24 and 48 hours. Target DLBCL cells were transduced via spinfection in the presence of 8 μg/mL polybrene (Sigma Aldrich), and incubated overnight. GFP-positive cells were sorted by fluorescence-activated cell sorting (FACS) using a Sony SH800S cell sorter (Sony Biotechnology, San Jose, CA), and seeded at 1 or 10 cells per well into 96-well plates. Clonal expansion was monitored, and screened for AID knockout by immunoblot analysis.

Knockdown of *AICDA* was achieved using the pLKO-3xLacO-mCherry lentiviral vector encoding a short hairpin RNA (shRNA) targeting *AICDA*. The shRNA sequence used was: 5’GTACC<u>CATTTCGTACTTTGGGACTTT</u>CTCGAG<u>AAAGTCCCAAAGTACGAAATG</u>TTTTTG3’. Lentiviral particles were produced and DLBCL cells transduced as described above. The top 10% of mCherry-positive cells were sorted by FACS to establish polyclonal knockdown cultures. Knockdown was confirmed after 3 days of incubation with IPTG using immunoblot analysis.

For AID overexpression, a retroviral expression construct; *LZRS-FLAG-AID-ER-IRES-ΔNGFR*, was generated by subcloning a codon-optimized and CRISPR/Cas9-resistant variant of mouse *Aicda* fused to the estrogen receptor alpha ligand-binding domain (AID-ERα) into the *LZRS-FLAG-IRES-ΔNGFR* backbone. Retroviral particles were produced by transfecting Phoenix-GaLV packaging cells with the retroviral construct using Genius transfection reagent, following the manufacturer’s instructions. Viral supernatants were harvested and centrifuged onto Retronectin-coated 24-well plates (Takara Bio Inc., Kusatsu, Japan) as per the manufacturer’s protocol. After removal of viral supernatant, DLBCL cells were added to the pre-coated wells and incubated for 3 days in IMDM supplemented with 20% FCS. Following transduction, cells were expanded and sorted for *ΔNGFR* expression by FACS, selecting the top 10% of ΔNGFR-positive cells using anti-NGFR antibody (ME20.4-1.H4, Miltenyi Biotec, Bergisch Gladbach, Germany). Polyclonal populations were verified for AID-ERα protein expression via immunoblotting. To confirm functional nuclear localization of AID-ERα upon 4-hydroxytamoxifen treatment, and surface IgM expression was assessed by flow cytometry following antibody staining, as an indicator of AID-driven class-switch-like activity.

### IgM staining

To assess IgM loss after nuclear translocation of AID-ERα clones, cells were seeded at 0.3 × 10^6^ cells/ml and cultured for 11 days in the presence 4-hydroxytamoxifen (4-OHT; Selleckchem, Houston, TX) at a concentration of 3 µM (HBL1), 0.3 µM (TMD8) or 1 µM (SU-DH-L6 and OCI-LY18). Cells were harvested at day 5 and day 11 of treatment, washed with FACS buffer (PBS, 0.5% BSA, 1 mM EDTA, 0.05% Sodium Azide), and stained with anti-IgM-PE (Southern Biotech, Birmingham, AL) (1:10000 for HBL1 and 1:2000 for TMD8) for 30 minutes at 4°C. Cells were washed and surface IgM expression was assessed by FACS, on a FACS CANTO II flow cytometer (BD Biosciences, Franklin Lakes, NJ). Data was analyzed using FlowJo (Tree Star Inc, Ashland, OR).

### *In vitro* competition assay

To assess the effect of AID knockdown on proliferation, mCherry-positive *AICDA*-shRNA clones were mixed with mCherry-negative WT cells at a 1:1 ratio. Cells were seeded at 0.3 × 10^6^ cells/ml and maintained in 10 mM isopropyl β-D-1-thiogalactopyranoside (IPTG, Sigma Aldrich) to induce shRNA expression, for the duration of the experiment. Cells were passaged every 3 days in the presence of IPTG. FACS flow cytometry was performed every three days to establish changes in population ratios using the BD FACS Symphony. Analysis was performed using FlowJo. The percent mCherry-positive cells were normalized to the initial seeding ratio with the WT competitor cells. Experiments were performed in triplicate.

### Cell cycle analysis

To evaluate cell cycle distribution, shRNA-expressing TMD8 cells were seeded at a density of 0.3 × 10⁶ cells/mL and cultured for 12 days in the presence or absence IPTG, with passaging every three days to maintain optimal growth conditions. At the end of the culture period, cells were pulse-labeled with 20 μM bromodeoxyuridine (BrdU; Sigma-Aldrich) for 1 hour at 37°C. Following BrdU incorporation, cells were washed with ice-cold phosphate-buffered saline (PBS) and fixed in 75% ethanol (pre-chilled at −20°C) for 1 hour. Fixed cells were subsequently permeabilized and partially digested by treatment with 0.4 mg/mL pepsin (Sigma-Aldrich) in 0.2 mM HCl for 25 minutes at 37°C. DNA denaturation was carried out by incubating cells in 2 M HCl for 25 minutes at 37°C, after which they were washed sequentially in 0.5% bovine serum albumin (BSA) in PBS (PBS-B), followed by 0.5% Tween-20 in PBS-B (PBS-TB). Cells were then stained with FITC-conjugated anti-BrdU antibody (clone B44; BD Biosciences) for 30 minutes at room temperature in the dark. After additional washing steps in PBS-B and PBS-TB, cells were stained with 0.1 μM TO-PRO-3 iodide (Invitrogen Life Technologies, Carlsbad, CA) in the presence of 500 μg/mL RNase A (Sigma-Aldrich) in PBS containing 0.5% BSA and 0.02% sodium azide. DNA content and BrdU incorporation were analyzed by flow cytometry.

### Immunoblotting

Cells were harvested, washed with ice-cold phosphate-buffered saline (PBS), and lysed in ice-cold radioimmunoprecipitation assay (RIPA) buffer composed of 50 mM Tris-HCl (pH 7.4), 150 mM NaCl, 1% Nonidet P-40, 0.5% sodium deoxycholate, and 0.1% sodium dodecyl sulfate (SDS). The lysis buffer was supplemented with EDTA-free protease inhibitor cocktail and phosphatase inhibitor tablets (PhosSTOP; both from Roche Diagnostics, Rotkreuz, Switzerland). Lysates were homogenized by brief vortexing and subsequently clarified by centrifugation. Protein concentrations were quantified using the bicinchoninic acid (BCA) assay (Sigma-Aldrich, St. Louis, MO, USA). Equal amounts of protein (15 μg per sample) were resolved on Precise™ 4–20% gradient Tris-SDS polyacrylamide gels (Thermo Fisher Scientific, Waltham, MA, USA) and transferred onto polyvinylidene difluoride (PVDF) membranes (Immobilon-P; EMD Millipore, Burlington, MA, USA). Membranes were blocked for 1 hour at room temperature (RT) in tris-buffered saline with 0.1% Tween-20 (TBS-T) supplemented with 5% non-fat dry milk. Primary antibodies were applied overnight at 4°C. After washing with TBS-T, membranes were incubated for 1 hour at RT with horseradish peroxidase (HRP)-conjugated secondary antibodies (goat anti-rabbit HRP or rabbit anti-mouse HRP; DAKO, Agilent Technologies, Santa Clara, CA, USA). Signal detection was performed using the Amersham™ ECL Prime chemiluminescent substrate (GE Healthcare BioSciences AB, Uppsala, Sweden), and bands were visualized with an appropriate imaging system. Quantification of protein band intensities was carried out using ImageJ software (imagej.net). The following primary antibodies were used: anti-AID clone 94.16 (kind gift from Prof. Hans-Martin Jäck, University of Erlangen-Nürnberg, Germany), NF-κB p105/p50 (Cell Signaling Technology, Danvers, MA, USA), BCL6 (DVI2V) XP (Cell Signaling Technology), IRF4 (E8H3S) XP (Cell Signaling Technology), MYC clone Y69 (Epitomics, Burlingame, CA, USA), and β-actin (clone C4; Millipore, Burlington, MA, USA).

### RNA-sequencing

Cells were seeded at 0.3 × 10^6^ cells /ml and passaged every 3 days. AID knockout cells and NT control cells were cultured for 2 weeks. AID-ER clones were treated for 11 days with tamoxifen at a concentration of 3 µM (HBL1), 0.3 µM (TMD8) or 1 µM (SU-DH-L6 and OCI-LY18). Total RNA was harvested using Tri Reagent (Merck KGaA, Darmstadt, Germany) extraction followed by purification using RNase minikit (Qiagen, Venlo, The Netherlands). 1 µg of RNA was used for sequencing library preparation, using TruSeq stranded mRNA library kit (Illumina, San Diego, CA), and sequenced paired-end on a Illumina HiSeq 4000, read length 100 bp. Sequence reads were pre-processed in Galaxy (https://usegalaxy.org), trimmed with Trim Galore and aligned with HISAT2 to hg38. Counts per gene were collected using Featurecounts and were uploaded in R2 Genomics Analysis and Visualization Platform (https://r2.amc.nl) for normalization, DESEq2 analysis and PCA analysis. R-Studio was used for visualization and heatmaps were generated by applying z-score normalization to the data. Gene set enrichment analyses (GSEA) and leading-edge (LE) analyses were performed using the GSEA software suite (http://www.broad.mit.edu/gsea) in combination with the Molecular Signature Database (MSigDB). Venn diagram–based gene overlap analysis was restricted to protein-coding genes.

Publicly available gene expression profile (GEP) datasets GSE31312 (39) and GSE117556 (40) were obtained from the NCBI Gene Expression Omnibus database. GEP datasets were analyzed by stratifying samples into activated B-cell-like (ABC), germinal center B-cell-like (GCB), and unclassified (UNC) subtypes. Within each subtype, patients were further categorized based on expression levels of *AICDA*. Specifically, individuals in the upper and lower quartiles of *AICDA* expression were selected for comparison. Differential gene expression analysis was performed using DESeq2, followed by GSEA and LE analysis. Primary sequencing data are publicly available through GEO accession number GSE308040.

### Statistics

The GraphPad Prism software package was used for statistical analyses of the cell culture and flow cytometry experiments (GraphPad Software, La Jolla, CA).

## RESULTS

### AID Regulates MYC and E2F Target Gene Expression In ABC-type DLBCL

To examine the effects of AID on global gene expression in ABC-type DLBCL, AID-deficient clones were generated by CRISPR-Cas9 gene targeting in the AID-expressing cell lines HBL1, TMD8, and OCI-LY10, and efficient knockout was confirmed by immunoblot analysis **(Figure 1A-C)**. Differential expression analysis of RNA sequencing data from three AID-deficient versus non-targeting (NT) guide RNA-expressing clones per cell line (DEseq2, FDR <0.1, log(FC) <-1 and >1) identified 186, 100, and 12 differentially expressed genes in HBL1, TMD8, and OCI-LY10, respectively **(Figure 1D-F)**, indicating that AID influences gene expression in these cell lines. Limited overlap between differentially expressed genes across the three lines indicated that AID’s transcriptional effects are context-dependent, with *AICDA*, encoding AID, as the only consistently differentially expressed gene **(Supplementary Figure 1A)**.

**Figure 1.**
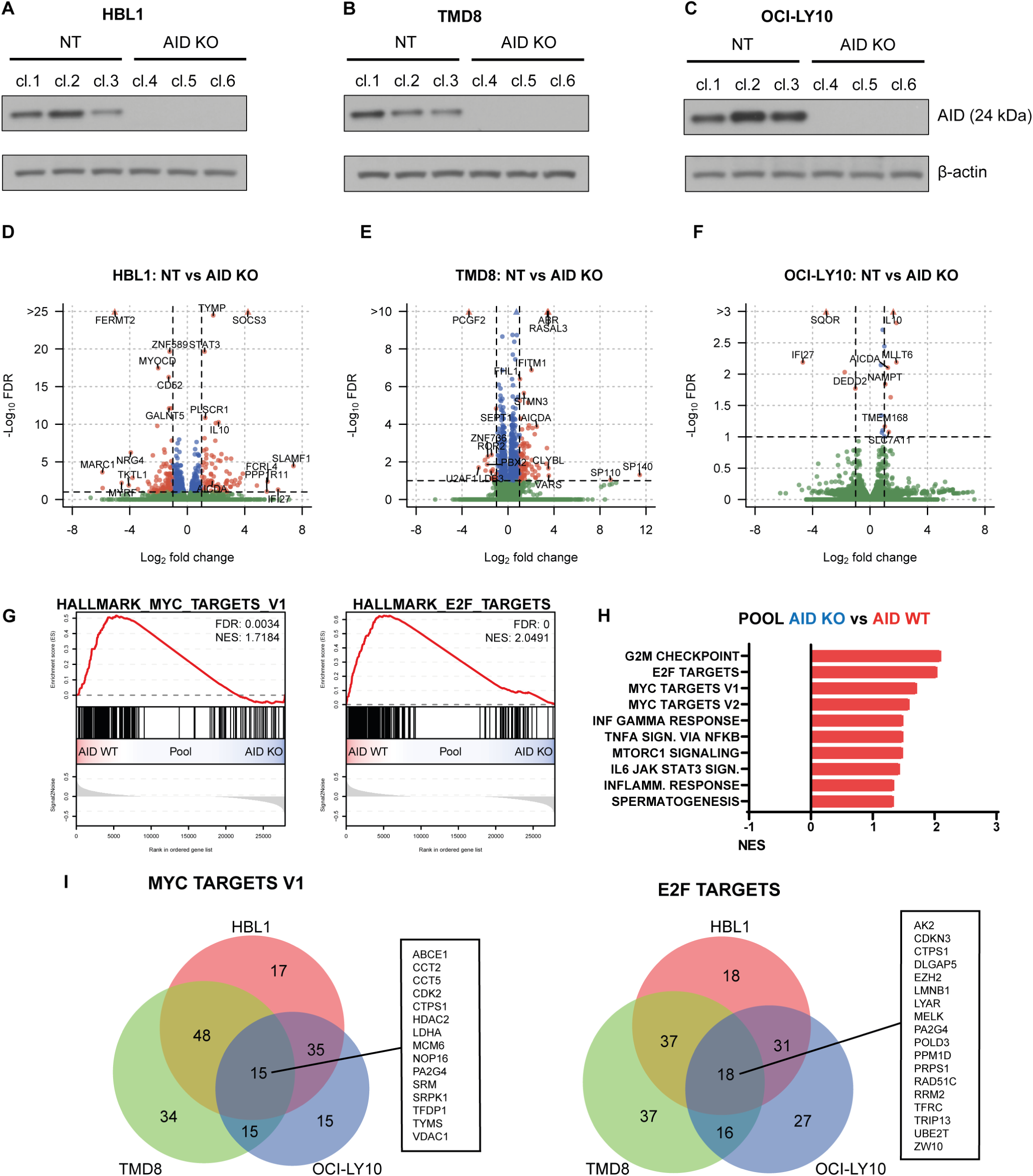
AID loss affects gene expression in ABC-type DLBCL and reduces MYC and E2F pathway activity. **A-C**, Immunoblot analysis of AID expression in CRISPR knockout (KO) and non-targeting (NT) control cells of HBL1 (**A**), TMD8 (**B**), and OCI-LY10 (**C**) cell clones. β-actin was used as a loading control. **D-F**, Volcano plots showing DESeq2 differential gene expression analysis on three independent CRISPR KO clones and corresponding NT control clones for each cell line: HBL1 (**D**), TMD8 (**E**) and OCI-LY10 (**F**). Significance was defined as false discovery rate (FDR) <0.1 and log fold change (FC) <-1 and >1. **G**, Pooled gene Set enrichment analysis (GSEA) plots depicting enrichment of MYC and E2F target genes in TMD8, HBL1 and OCI-LY10 CRISPR KO and NT clones. FDR and normalized enrichment score (NES) values are shown in the plots. **H**, Bar graph showing enriched pathways identified by gene set enrichment analysis in the pooled cell clones. Gene sets with a FDR < 0.2 are displayed. Gene sets significantly enriched in AID WT clones are shown in red; no gene sets met significance criteria in AID KO clones. The x-axis represents enrichment scores. **I**, Venn diagrams illustrating the overlap of leading-edge genes from the enriched gene sets.

To obtain a broader view of pathways affected by AID, we performed gene set enrichment analysis (GSEA) on pooled RNA sequencing data from HBL1, TMD8, and OCI-LY10 AID-deficient versus AID-proficient clones. The top enriched gene sets in the AID-proficient pool were G2M checkpoint, E2F targets, and MYC targets (FDR<0.2) **(Figure 1G-H)**. MYC and E2F targets represent key oncogenic pathways associated with tumor cell growth and aggressive disease, and enrichment of G2M checkpoint genes suggests enhanced cell cycle progression. MYC target genes were consistently enriched in AID-positive clones of all three lines, and E2F target genes were significantly enriched in two of three lines (FDR<0.2) **(Supplementary Figure 1B-D)**. No gene sets were significantly enriched in the AID-deficient clones **(Figure 1H)**. Leading-edge analysis (LEA) identified 15 MYC and 18 E2F leading-edge genes shared between the three cell lines **(Figure 1I, Supplementary Table 1)**. *CTPS1* and *PA2G4* were present in both MYC and E2F leading-edge sets and encode a rate-limiting enzyme in *de novo* pyrimidine synthesis and an RNA-binding protein involved in growth regulation, respectively. Together, these data indicate that AID expression is associated with activation of MYC and E2F target genes and may promote proliferation and cell cycle progression in ABC-type DLBCL.

### Complementation of AID expression in AID-deficient ABC-type DLBCL Restores MYC and E2F Target Gene Expression

To further establish the functional link between AID expression and MYC/E2F pathway activity, AID-deficient TMD8 (n=3) and HBL1 (n=3) clones were transduced with a CRISPR/Cas9-resistant, inducible AID-ERα fusion construct allowing nuclear AID expression upon tamoxifen treatment, or a ΔNGFR control vector **(Figure 2A-B)**. Canonical AID activity at the immunoglobulin heavy chain (*IGH*) locus was assessed by monitoring surface IgM expression as a surrogate readout of SHM/CSR-induced *IGH* disruption (37, 38). After five days of tamoxifen treatment, a subpopulation of AID-ERα-expressing cells showed marked loss of IgM surface expression **(Supplementary Figure 2)**, indicating effective targeting of *IGH*. TMD8 cells with nuclear AID-ERα displayed a 12.7-fold increase in IgM-negative cells compared to untreated controls (n=3), and HBL1 cells showed a 3.7-fold increase (n=3), confirming robust functional activity of the AID-ERα fusion.

**Figure 2.**
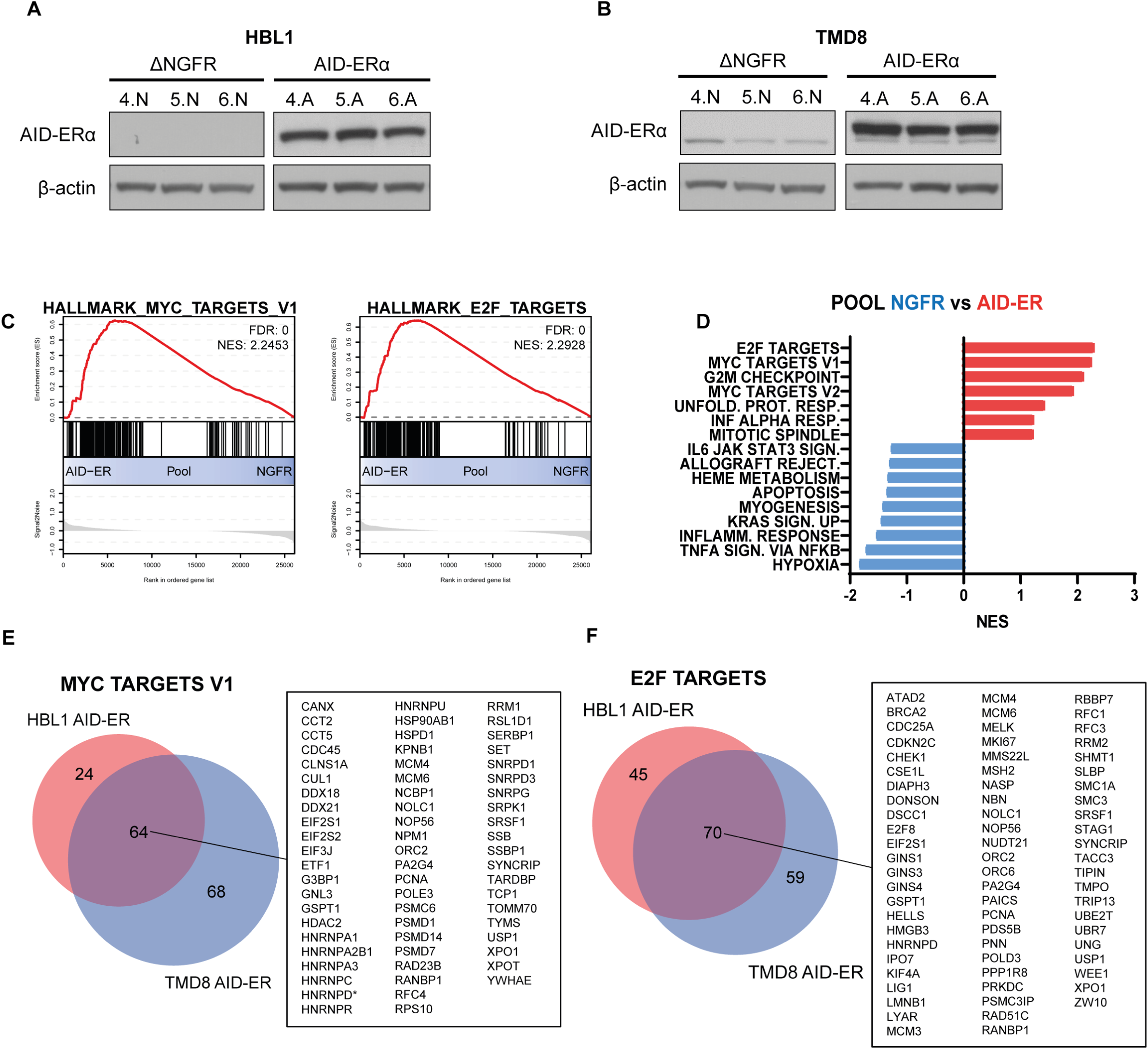
Overexpression of AID in AID deficient ABC-type DLBCL reverses MYC and E2F pathway activity. **A-B**, Immunoblot analysis showing AID-ERα expression in three independent AID knockout clones of HBL1 (**A**) and TMD8 (**B**) cells transduced with either the AID-Erα fusion construct or the ΔNGFR control construct. β-actin was used as a loading control. **C**, GSEA plots depicting enrichment of MYC and E2F target gene sets in pooled independent TMD8 and HBL1 AID-ERα and ΔNGFR clones. FDR and NES values are shown in the plots. **D**, Bar graph showing enriched pathways identified by pooled gene set enrichment analysis in AID-ERα and ΔNGFR cell clones. Gene sets with a false discovery rate (FDR) < 0.2 are displayed, with AID-ERα–enriched gene sets shown in red and ΔNGFR-enriched gene sets in blue. The x-axis indicates enrichment scores. **E-F**, Venn diagrams illustrating the overlap of leading-edge (LE) genes from the MYC and E2F target gene sets across HBL1 and TMD8 clones overexpressing AID-ERα.

To capture transcriptional consequences of sustained AID activity, tamoxifen treatment was extended to 11 days. RNA sequencing then revealed 1 and 16 differentially expressed genes in AID-ERα–proficient versus AID-deficient control cells for TMD8 and HBL1, respectively (FDR <0.1, log2(FC) <-1 and >1, data not shown). GSEA of combined TMD8 and HBL1 datasets demonstrated significant enrichment of E2F and MYC target gene sets, as well as G2M checkpoint gene set, in AID-ERα–expressing cells **(Figure 2C-D)**, indicating that AID promotes cell cycle progression and proliferation-associated pathways. Comparison of leading-edge genes showed that 64 MYC pathway genes and 70 E2F pathway genes enriched in TMD8 AID-ERα cells were also enriched in HBL1 AID-ERα cells **(Figure 2E-F)**, demonstrating a conserved effect. Shared upregulated genes included DNA replication genes (*CDC45, MCM4, MCM6, NMP1, PCNA, TYMS, RRM1, RCF4, ORC4*) and key cell cycle regulators (*ATAD2, BRCA2, CDC25A, CDKN2C, CHEK1, E2F8, GINS1, GINS2, GINS4, HELLS, KIF4A, LIG1, LMNB1*). Thus, reintroduction of AID into AID-deficient cells restores MYC and E2F target gene expression and reverses the transcriptional consequences of AID loss, supporting a direct role for AID in enhancing MYC/E2F pathway activity.

### Inhibition of AID Reduces Cell Cycle Progression and Proliferation in ABC-type DLBCL

Initial comparisons of CRISPR/Cas9-derived AID-deficient and AID-proficient ABC-type DLBCL clones did not reveal significant differences in proliferation or cell cycle distribution, likely due to clonal variation (data not shown). To circumvent clonal artifacts, TMD8 cells were transduced in bulk with inducible short hairpin RNA (shRNA) targeting *AICDA* or a non-targeting (NT) control shRNA. IPTG induction of the AID-targeting shRNA reduced AID protein expression by 52% after 3 days, while the NT shRNA did not impact AID expression **(Figure 3A)**.

**Figure 3.**
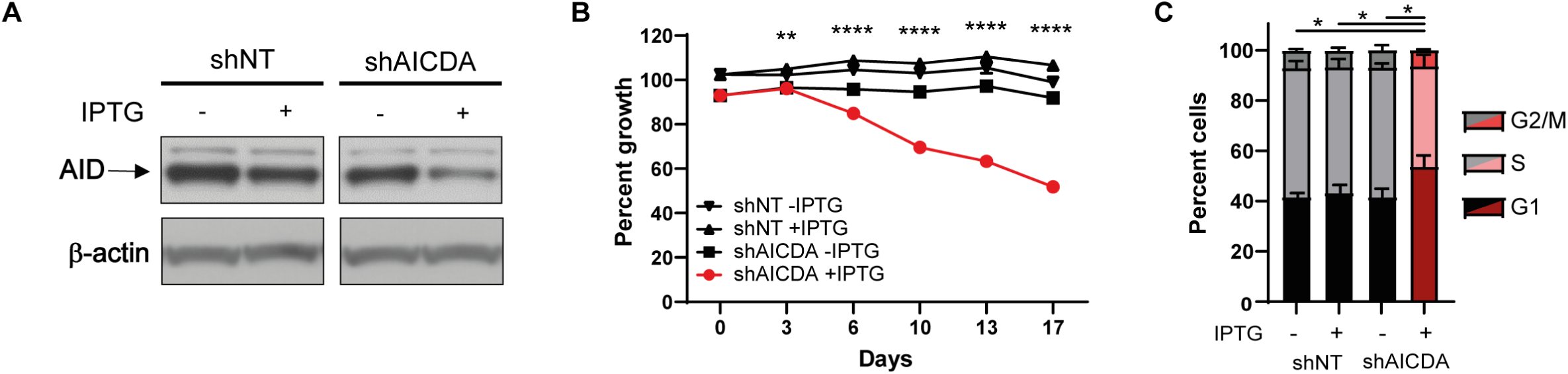
Knockdown of AID in TMD8 cells affects proliferation. **A**, Immunoblot analysis of AID knockdown in shAICDA and shNT control cells following 3 days of IPTG treatment. β-actin served as a loading control. **B**, Competition assay in which mCherry-positive shAICDA and shNT clones were mixed with non-fluorescent wild-type (WT) cells in biological triplicates and treated with 10 mM IPTG to induce shRNA expression. Cell proliferation was assessed by flow cytometry (FACS), cell counts were normalized to the initial WT-to-shRNA seeding ratio. Statistical significance was determined by one-way ANOVA with Bonferroni correction, comparing shAICDA + IPTG (red circles) to shAICDA – IPTG (black squares) at each time point. **C**, Cell cycle distribution after 12 days of IPTG treatment, assessed by BrdU incorporation and flow cytometry. Biological triplicates are depicted. Cells in G1 phase (black/dark red), S phase (light grey/pink), and G2/M phase (grey/red) are depicted. One-way ANOVA with Bonferroni correction was performed (* p<0.05; ** p<0.01; **** p<0.0001).

To assess the impact of AID inhibition on proliferation, an *in vitro* competition assay was performed in which fluorescently labeled AID-knockdown or NT control cells **(Supplementary Figure 3A)** were mixed 1:1 with parental TMD8 cells and cultured for 17 days with or without IPTG. This revealed that AID inhibition conferred a pronounced proliferative disadvantage, evident from day 3 of IPTG treatment, ultimately resulting in a 51.4% reduction in AID-knockdown cells relative to IPTG induced shNT control cells after 17 days. In contrast, non-IPTG induced shAICDA and shNT shRNA-expressing cells proliferated comparably **(Figure 3B)**.

To determine whether reduced cell numbers was due to apoptosis, cultures were treated with the pan-caspase inhibitor QVD-OPH, which failed to rescue the proliferative defect **(Supplementary Figure 3B)**, indicating that AID inhibition did not induce apoptosis. BrdU incorporation assays showed that AID inhibition significantly increased the proportion of cells in G1 phase from 42% to 54% (p<0.05), with a corresponding decrease in S phase cells (p<0.05) **(Figure 3C)**. Relative to NT shRNA controls with or without IPTG, AID knockdown increased the G1 fraction by 10% and 12%, respectively, accompanied by a decrease in S phase cells (p<0.05). These findings demonstrate that AID inhibition impairs G1/S transition and proliferation, indicating that AID expression promotes cell cycle progression in ABC-type DLBCL, in line with its association with MYC and E2F pathway activity.

### AID Expression Correlates with MYC and E2F Pathway Activation in DLBCL Patient Tumors

Having established a causal link between AID expression and MYC/E2F pathway activity in ABC-type DLBCL cell lines, we next investigated whether AID expression is similarly associated with these pathways in primary DLBCL tumors. Two publicly available gene expression profiling datasets were analyzed: the Visco *et al.* cohort (n=498) of newly diagnosed DLBCL patients treated with R-CHOP (39), and the Sha *et al.* cohort (n=928), which evaluated the addition of bortezomib to R-CHOP (40).

In each dataset, tumors were stratified by ABC, GCB, or unclassified (UNC) subtype and further subdivided by *AICDA* mRNA expression (upper vs lower quartile). Differential expression analyses revealed distinct AID-associated transcriptional profiles. In the Visco *et al.* cohort, 143 and 14 and 1 genes were differentially expressed in ABC, GCB and UNC subtypes, respectively, while in the Sha *et al.* cohort, 203 and 20 genes were differentially expressed in ABC and GCB subtypes, respectively **(Figure 4A, Supplementary Figure 4A-B)**. Few genes were consistently differentially expressed across subtypes within each dataset (one gene in Visco *et al.*, and two in Sha *et al.*), indicating that AID’s impact on individual genes is largely subtype-dependent.

**Figure 4.**
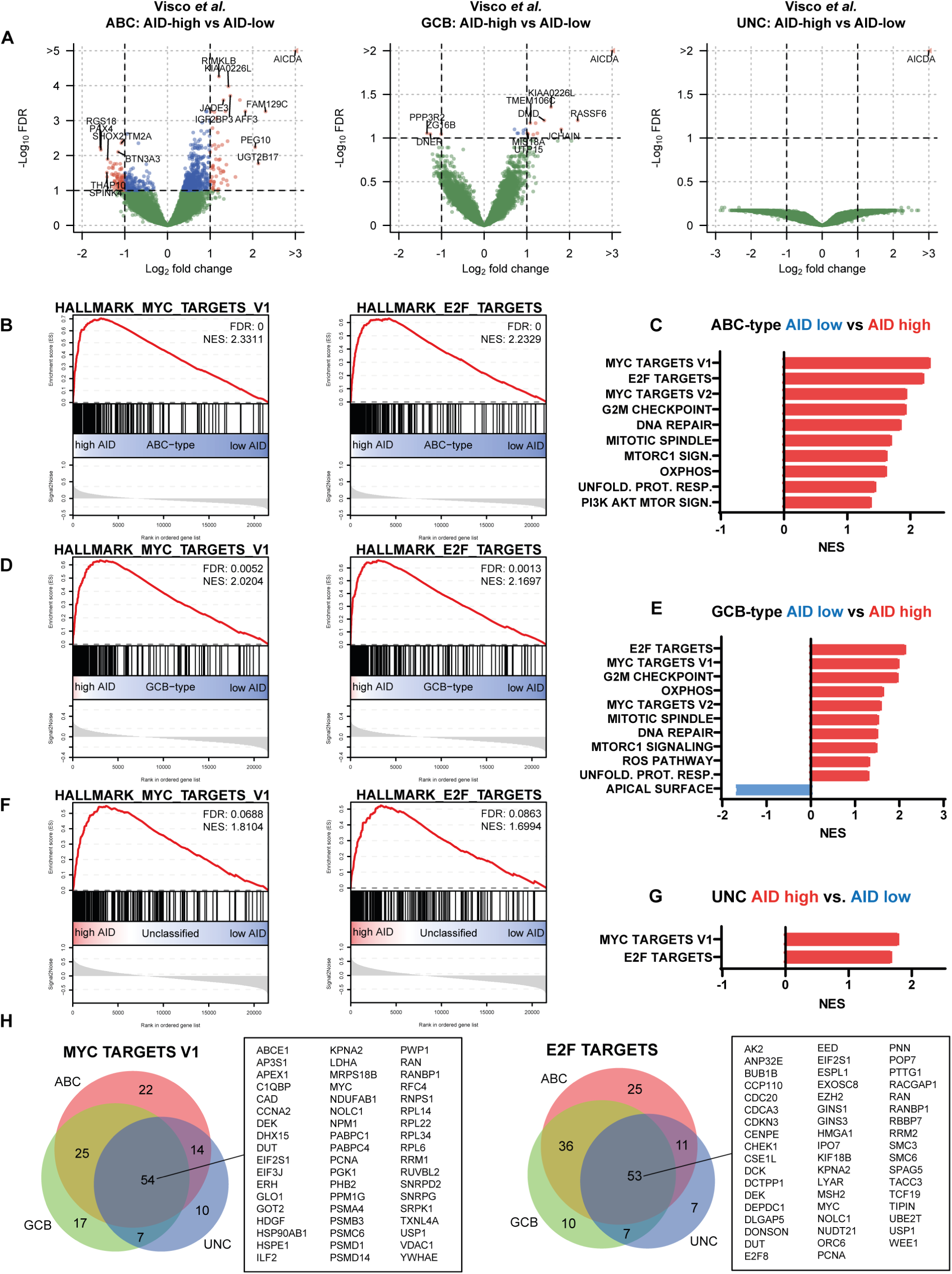
DLBCL patients with high AID expression show enrichment of MYC and E2F transcriptional programs. **A**, Volcano plots showing differential gene expression analysis on ABC-type (left panel), GCB-type (middle panel) and unclassified (right panel) DLBCL patient samples from Visco *et al.* comparing the upper and lower quartiles of AID expression. Significance was defined as FDR <0.1 and log(FC) <-1 and >1. **B-G**, GSEA enrichment plots for MYC and E2F signatures in ABC-type (**B**), GCB-type (**D**), and unclassified (**F**) DLBCL patient samples. FDR and NES values are shown in the plots. Significantly enriched pathways between AID-high (red) and AID-low (blue) expressing groups (FDR < 0.2) are shown for ABC-type (**C**), GBC-type (**E**), and unclassified (**G**). **H**, Venn diagrams illustrating the overlap of leading-edge genes of the MYC (left panel) and E2F (right panel) enriched gene sets across the DLBCL subtypes.

Despite limited overlap at the single-gene level, GSEA revealed consistent enrichment of MYC and E2F target genes in AID-high tumors across all subtypes in the Visco *et al.* cohort (FDR<0.2; **Figure 4B-G**). In the Sha *et al.* cohort, MYC and E2F target genes were significantly enriched in AID-high ABC tumors (FDR<0.2), and GCB tumors showed a non-significant trend toward MYC target gene enrichment (FDR<0.25) **(Supplementary Figure 5A-D)**. Additional gene sets significantly enriched in AID-high tumors included MTORC1 signaling, unfolded protein response, G2M checkpoint, and mitotic spindle gene sets **(Figure 4C, 4E, Supplementary Figure 5A-D)**.

Leading-edge analyses demonstrated extensive overlap of MYC and E2F leading-edge genes across DLBCL subtypes within each dataset. In the Visco *et al.* cohort, 54 MYC and 53 E2F leading-edge genes were shared across ABC, GCB, and UNC subtypes **(Figure 4H)**. In the Sha *et al.* dataset, 75 MYC and 77 E2F leading-edge genes were shared between ABC and GCB subtypes **(Supplementary Figure 5E)**. Cross-cohort comparison showed substantial overlap within each subtype (56 and 61 overlapping MYC and E2F genes in ABC, and 64 and 54 in GCB, respectively; **Supplementary Figure 5E**). Collectively, these findings indicate that higher AID expression in DLBCL patients is robustly associated with increased MYC and E2F pathway activity across molecular subtypes. While this correlation, together with the *in vitro* data, aligns with reports linking AID expression to poorer outcomes in DLBCL (24, 25), the independent prognostic value of AID remains unresolved and additional studies are required to disentangle AID-specific effects from other clinical and molecular risk factors.

### AID Expression Skews Cell-of-Origin Gene Expression Programs in DLBCL

Given the association between AID and cell cycle progression, we next investigated whether AID also affects cell-of-origin (COO) gene expression programs that define ABC and GCB DLBCL. We first compared AID-proficient and AID-deficient ABC-type cell lines. In HBL1, GSEA revealed significant enrichment of GCB stratification genes in AID-deficient cells (FDR<0.2) and a non-significant trend toward enrichment of ABC-type stratification genes in AID-proficient cells **(Figure 5A)**. AID-deficient TMD8 and OCI-LY10 cells also showed non-significant enrichment of ABC-stratification genes in AID-proficient cells (data not shown), without GCB gene enrichment in these lines. Leading-edge analysis identified *IRF4*, *PTPN1*, and *SLA* as ABC stratification genes consistently enriched in AID-proficient cell lines **(Figure 5B)**. IRF4 is a key driver of the ABC subtype (9), and PTPN1 and SLA are tumor suppressors that negatively regulate NF-κB in an indirect fashion (41, 42). No GCB stratification genes were commonly enriched across AID-deficient cell lines.

**Figure 5.**
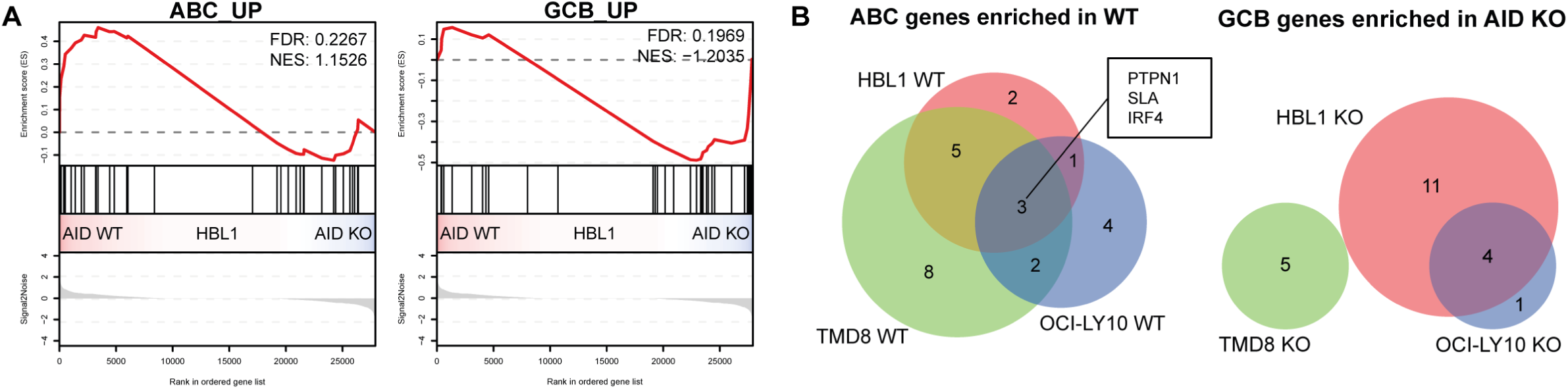
AID loss in ABC-type DLBCL cells skews gene expression profile to GCB-type DLBCL. **A**, GSEA enrichment plots of ABC (left panel) and GCB (right panel) stratification genes in the ABC-type DLBCL cell line HBL1 comparing three independent AID knockout (KO) and AID non-targeting (NT) clones. **B**, Venn diagrams showing the overlap of enriched ABC stratification genes in NT clones (left panel) and enriched GCB stratification genes in AID KO clones (right panel) identified by leading-edge analysis across the cell clones of HBL1, TMD8, and OCI-LY10.

To test whether AID can shift GCB-type DLBCL cell lines toward an ABC-like profile, we overexpressed AID in GCB-type SU-DHL-6 and OCI-LY18 cells that express endogenous AID at low levels. Cells were transduced with inducible AID-ERα and treated with tamoxifen to induce nuclear localization, and AID enzymatic activity was confirmed by loss of surface IgM **(Supplementary Figure 6)**. GSEA comparing AID-overexpressing versus control (ΔNGFR) cells showed significant enrichment of ABC-type stratification genes in AID-overexpressing lines (FDR<0.2) and enrichment of GCB-type stratification genes in controls (FDR<0.2) **(Figure 6A)**. Heatmap visualization of COO-associated genes in SU-DHL-6 highlighted this shift, with ABC-type stratification genes showing increased expression in AID-overexpressing cells, whereas GCB-type genes were underrepresented **(Figure 6B)**. In AID-overexpressing SU-DHL-6 and OCI-LY18 cells, 4 ABC-type genes were consistently enriched. In the ΔNGFR control SU-DHL-6 and OCI-LY18 cells, 10 GCB-type genes were found enriched **(Figure 6C)**.

**Figure 6.**
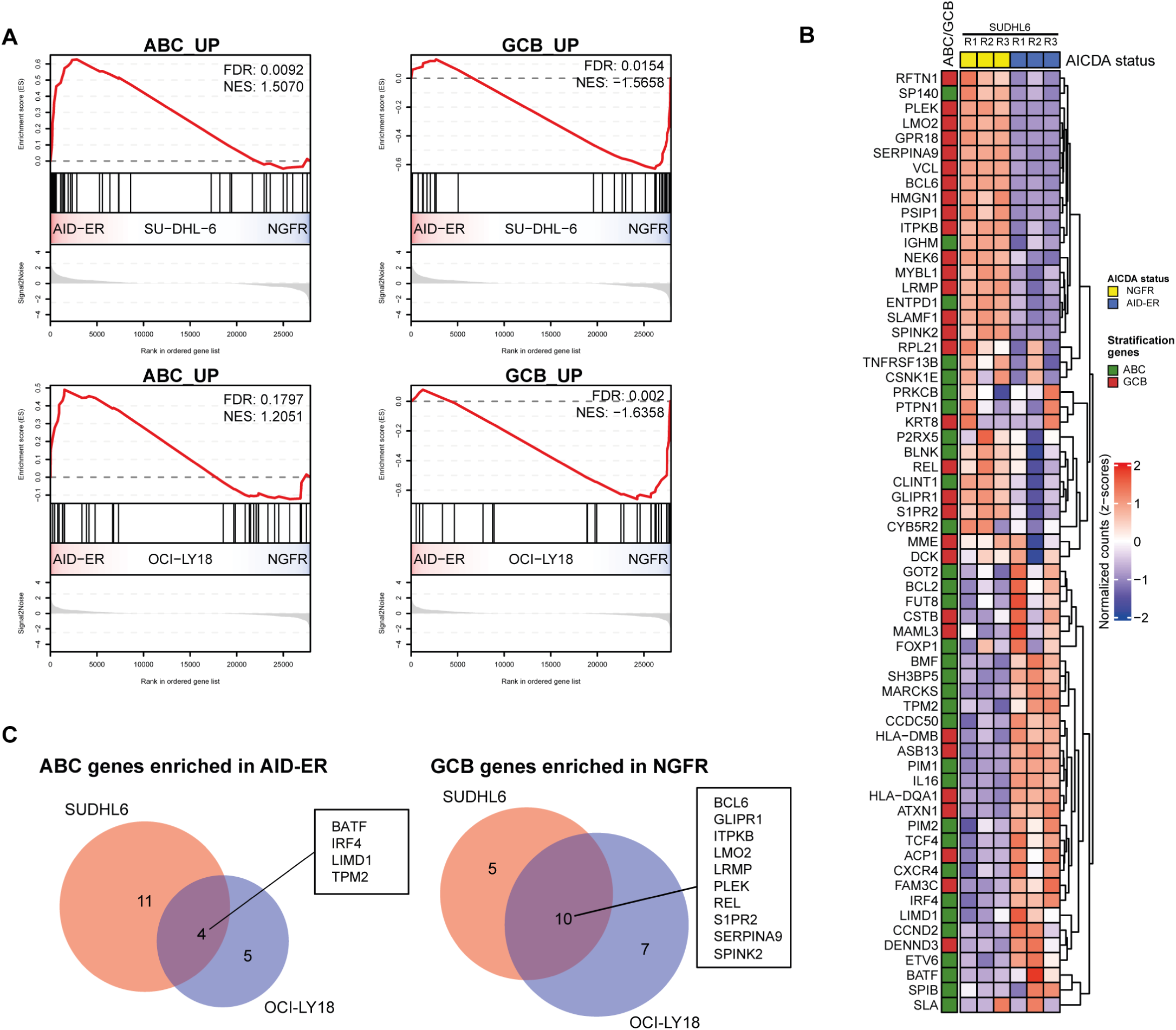
AID overexpression in GCB-type DLBCL skews gene expression profile to ABC-type DLBCL. **A**, GSEA of ABC (left panels) and GCB (right panels) stratification genes in biological triplicates of GCB-type DLBCL cell lines SU-DHL-6 (upper panels) and OCI-LY18 (lower panels) expressing AID-ERα or ΔNGFR control. FDR and NES values are indicated in the enrichment plots. **B**, Heatmap showing the z-score-normalized expression of ABC and GCB stratification genes in SU-DHL-6 cells. **C**, Venn diagram of ABC stratification genes enriched in AID-ERα expressing GCB-type DLBCL cell lines SU-DHL-6 and OCI-LY18 (left panel). Venn diagram of GCB stratification genes enriched in ΔNGFR control cell lines (right panel).

To further substantiate the ABC-type shift, we analyzed protein expression of COO regulators in AID-ERα-expressing SU-DHL-6 cells. AID overexpression led to a 2.5-fold increase in IRF4 protein, a master ABC regulator **(Figure 7A-B, 7I)**, while the key GCB regulator BCL6 showed a modest decrease **(Figure 7B, 7H)**. AID-overexpressing cells also displayed increased canonical NF-κB activity, evidenced by decreased p105 and increased p50 protein levels **(Figure 7C, 7F-G)**. Consistent with elevated MYC pathway activity in AID-expressing DLBCL, MYC protein levels were modestly increased (1.5-fold) in AID-overexpressing cells.

**Figure 7.**
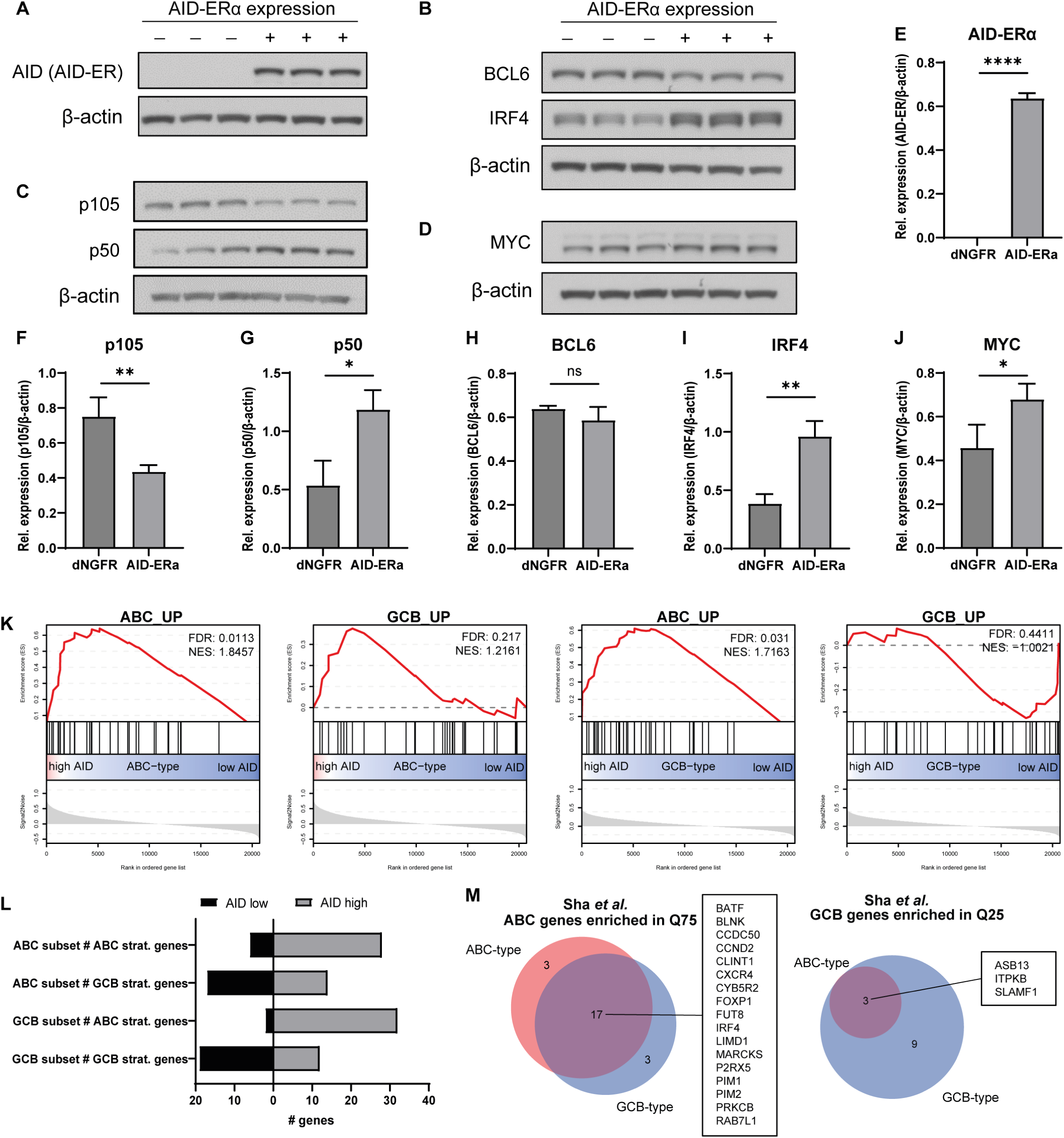
AID overexpression affects levels of DLBCL cell-of-origin (COO) proteins, and AID expression correlates with ABC stratification genes in DLBCL patients. **A–D**, Immunoblot analysis of AID-ERα (**A**), p105, p50 (**B**), BCL6, IRF4 (**C**) and MYC (**D**) protein expression in biological triplicates of the GCB-type DLBCL cell line SU-DHL-6 overexpressing AID-ERα or ΔNGFR control cells. Cells were seeded and maintained at 0.3 × 10^6 cells/ml with passaging every three days for 11 days in the presence of tamoxifen. **E–J**, Quantification of protein expression by densitometry of AID-ERα (**E**), p105 (**F**), p50 (**G**), BCL6 (**H**), IRF4 (**I**) and MYC (**J**). Statistical significance was assessed using a two-tailed unpaired *t*-test (* p<0.05; ** p<0.01; **** p<0.0001). **K**, Enrichment plots of ABC (first and third panels) and GCB (second and fourth panels) stratification gene sets from GSEA analyses of ABC-type (left two panels) and GCB-type (right two panels) DLBCL patients from Sha *et al.* Comparisons were performed between patients in the upper versus lower quartile of AID expression. **L**, Bar graphs showing the number of ABC and GCB stratification genes with higher expression in the upper quartile versus the lower quartile of AID expression (grey) or higher expressed in the lower quartile than the upper quartile of AID expression (black), stratified by ABC-type and GCB-type DLBCL patients from Sha *et al*. **M**, Venn diagrams depicting overlap in enriched genes: ABC stratification genes enriched in the AID upper quartile of both ABC-type and GCB-type DLBCL (left panel), and GCB stratification genes enriched in the AID lower quartile of both ABC-type and GCB-type DLBCL (right panel).

To extend these observations to primary tumors, we performed GSEA using COO gene sets on public datasets stratified by AID expression within each subtype. In the Sha *et al.,* cohort (40), ABC-type stratification genes were significantly enriched in AID-high tumors in both ABC and GCB subtypes (FDR<0.2), no enrichment of GCB genes was detected in AID-low ABC DLBCL **(Figure 7K)**. Most ABC-type stratification genes were expressed at higher levels in AID-high tumors irrespective of subtype (28 and 32 of 34 ABC genes in ABC-type and GCB-type tumors, respectively) **(Figure 7L)**. GCB stratification genes were more evenly distributed between AID-high and AID-low tumors, with a slight predominance in AID-low groups (17 and 19 of 31 GCB genes in ABC-type and GCB-type tumors, respectively). Similar patterns were observed in the Visco *et al.,* cohort (39), with more ABC-type genes upregulated in AID-high tumors and more GCB-type genes upregulated in AID-low tumors **(Supplementary Figure 7A)**. Principal component analysis revealed a pseudotime-like gradient along PC1 from GCB-type AID-low to GCB-type AID-high to ABC-type AID-low to ABC-type AID-high **(Supplementary Figure 7B)**. Leading-edge analysis in the Sha *et al.,* cohort identified 17 ABC stratification genes commonly enriched in AID-high (upper quartile: Q75) ABC and GCB tumors and 3 GCB stratification genes enriched in AID-low (lower quartile: Q25) ABC and GCB tumors **(Figure 7M)**. Notably, the master ABC regulator IRF4 and its coregulator BATF in ABC-type DLBCL (22, 43), were shared ABC leading-edge genes in AID-high tumors across subtypes.

Collectively, these findings indicate that AID not only maintains ABC stratification gene expression in ABC-type DLBCL cell lines but can also induce ABC-type programs in GCB-type cells. The increase in IRF4 expression and NF-κB activity in GCB cells with induced AID, together with skewing of COO signatures toward an ABC-like phenotype in AID-high patient tumors, supports a mechanistic role for AID in shaping DLBCL COO gene expression programs.

## DISCUSSION

AID is indispensable for adaptive immunity, enabling antibody diversification through SHM and CSR in germinal center B cells (1, 2). However, this genomic destabilizing activity drives B-cell lymphomagenesis through off-target DNA double-strand breaks at non-Ig loci (29, 32, 34, 44, 45), including MYC and BCL6, promoting translocations characteristic of Burkitt’s lymphoma and DLBCL (33, 46), and aberrant SHM fueling clonal evolution and treatment resistance (26, 47–49). Beyond these genomic effects, our findings reveal a broader role for AID in DLBCL pathogenesis, potentially linked to its deamination activity.

Our study shows that AID expression in ABC-type DLBCL drives MYC and E2F pathway activity, enhances proliferation, and reinforces ABC-type transcriptional programs. AID loss or inhibition impaired these pathways and delayed cell cycle progression, effects reversed by AID complementation. Similarly, AID-MYC/E2F correlations in primary DLBCL tumors indicate these effects extend beyond cell lines.

AID’s influence on gene expression may stem from deaminating 5-methylcytosine to initiate active DNA demethylation and epigenetic remodeling. Supporting evidence includes germinal center B cells from AID-deficient mice showing lost methylation diversity at SHM-targeted genes involved in B-cell development and proliferation (8). AID-overexpressing lymphomas exhibit increased methylation heterogeneity (36), and AID-TET2 cooperation regulates FANCA in DLBCL (50). AID knockdown inhibits proliferation and invasion in bladder cancer via methylation changes (51), alters hematopoietic stem cell differentiation with global hypermethylation (52), and is required for epithelial-mesenchymal transition (EMT) in breast cancer by demethylating transcription factor promoters (53). However, AID’s role in active demethylation remains controversial. AID-deficient hyper-IgM patients and mice show no demethylation at predicted hotspots (54), AID disfavors modified cytosines biochemically (55), and primordial germ cell demethylation attributed to AID may reflect passive replication or genetic confounders (5, 6, 56). Thus, while the effects of AID on DLBCL gene expression and subtype identity observed in this study align with locus-specific epigenetic regulation, its global role is context-dependent warranting further mechanistic studies.

Our data link AID to increased IRF4 protein and NF-κB1 activity (p105-to-p50 cleavage). IRF4, central to ABC-DLBCL identity and survival, cooperates with NF-κB to drive its gene program (57). NF-κB1 predominates in GCB-DLBCL, while NF-κB2 is associated with the ABC subtype. Modulation of these proteins shifts subtype identity, with p100 silencing inducing GCB-like states and p52/p100 promoting ABC-like phenotypes (58). AID-overexpression increasing NF-κB1 in GCB cells may thus influence subtype plasticity. Shared BATF enrichment in AID-high ABC/GCB tumors further supports this, as BATF partners with IRF4 at composite elements to drive ABC heterogeneity (43). As ABC-DLBCL depends on IRF4, and its downregulation impairs growth (57). Immunomodulatory drugs (IMiDs) like lenalidomide indirectly target IRF4 by ubiquitination involving the E3 ligase component cereblon (59, 60). Thus, modulating IRF4-BATF or NF-κB in AID-high, ABC-like DLBCL holds therapeutic promise.

In conclusion, AID regulates transcriptional programs linked to cell cycle progression, proliferation, and DLBCL subtype identity beyond mutagenesis, aligning with its multifaceted B-cell roles (35, 61–63). Since selective AID inhibitors are being developed (64, 65), and APOBEC3 targeting offers parallels (66–68), our findings warrant exploring AID/APOBEC blockade to disrupt oncogenic pathways and subtype plasticity in DLBCL.

## ACKNOWLEDGEMENTS

The authors thank the Guikema laboratory members for fruitful discussions, and the flow cytometry core facility at the Amsterdam UMC for technical assistance.

This work was funded by Cytura Therapeutics (Oss, the Netherlands), and supported by a PhD fellowship grant from the Amsterdam UMC to T.P.vD.

## AUTHORSHIP CONTRIBUTIONS

L.H.G. and T.P.vD. designed the study, performed experiments, analyzed data, and wrote the manuscript. M.F.dR. and G.dW. analyzed data and edited the manuscript. R.J.B., M.S., A.vG., and C.J.M.vN. edited the manuscript. J.E.J.G. designed and directed the study, analyzed data, and wrote the manuscript.

