## Supplementary Figures for "AID Shapes Proliferation and Cell-of-Origin-associated Transcriptional Programs in Diffuse Large B-cell Lymphoma"

Supplementary Figure 1

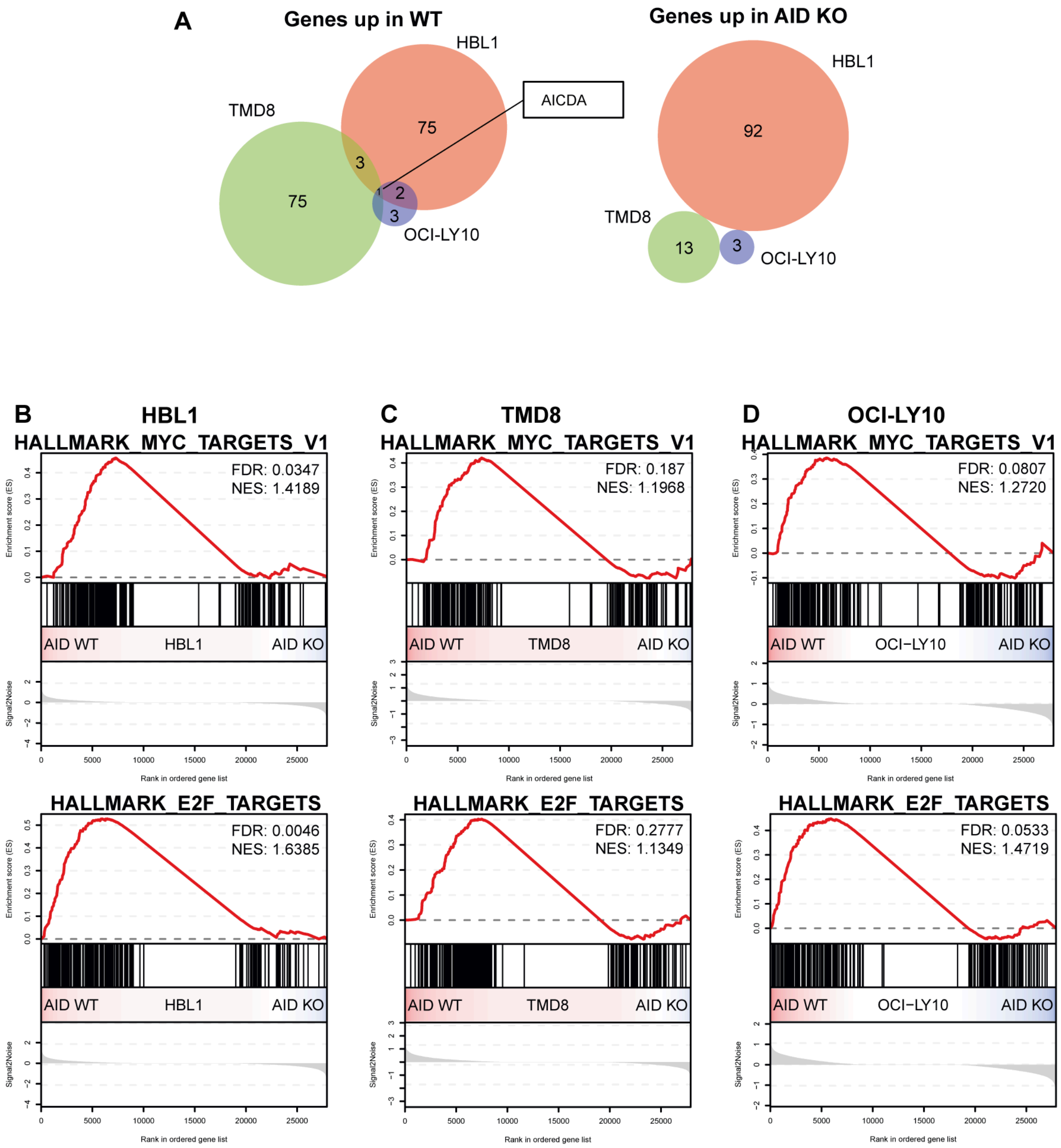


**Supplementary Figure 1. AID loss alters gene expression in ABC-type DLBCL cell lines and reduces MYC and E2F pathway activity across three independent models.**
**A**, Venn diagrams showing overlap of genes upregulated in NT (left panel) or AID KO (right panel) cells of HBL1, TMD8, and OCI-LY10, as identified by differential expression analysis. **B-D**, GSEA enrichment plots of MYC target genes (upper panels) and E2F target genes (lower panels) in HBL1 (**B**), TMD8 (**C**), and OCI-LY10 (**D**) CRISPR clones, based on three independent AID KO and NT clones per cell line.

Supplementary Figure 2

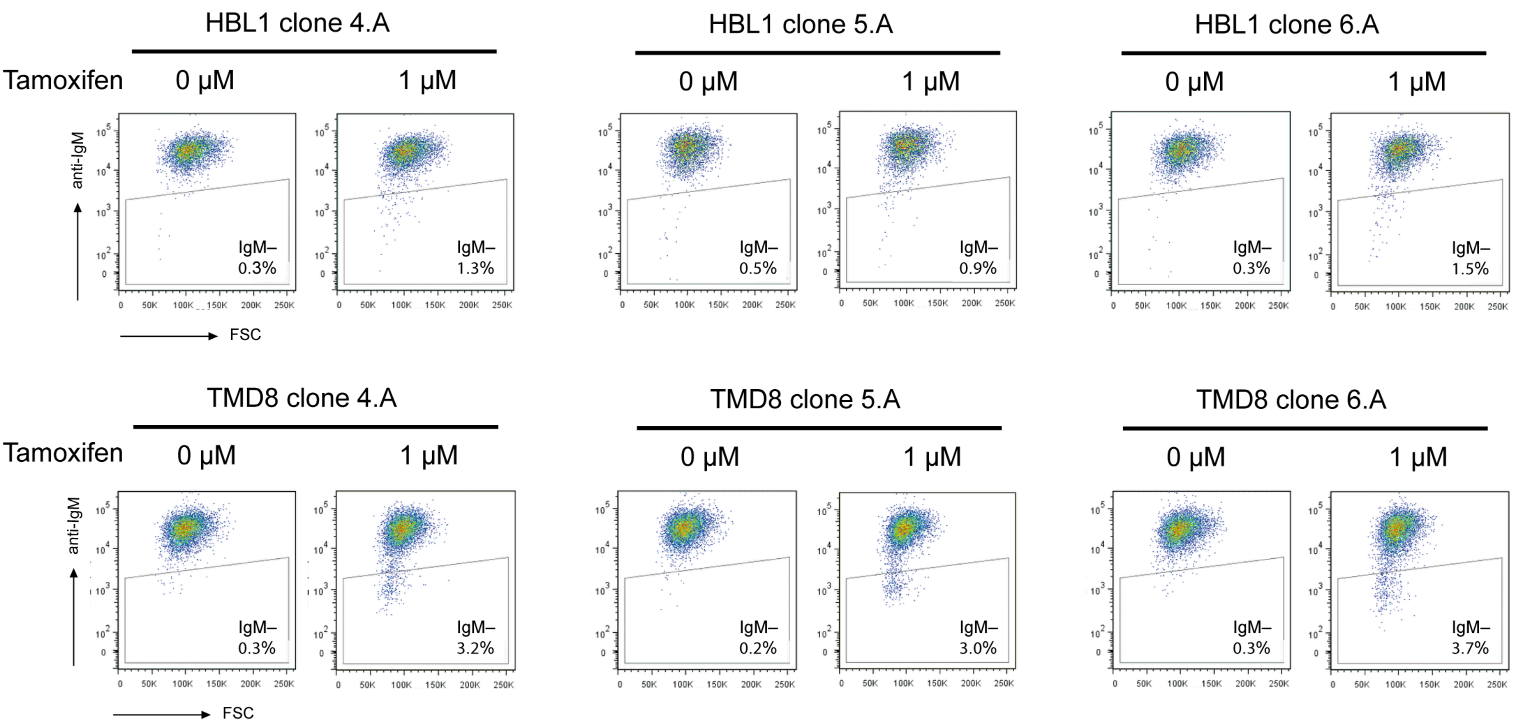


**Supplementary Figure 2. Assessment of AID-ERα activity upon tamoxifen treatment in ABC-type DLBCL AID KO clones HBL1 and TMD8.**
Flow cytometry plots showing surface IgM expression in independent HBL1 (upper panels) and TMD8 (lower panels) AID-ERα clones cultured for 5 days in the presence or absence of tamoxifen. Forward scatter (FSC) is shown on the x-axis and surface IgM expression on the y-axis.

Supplementary Figure 3

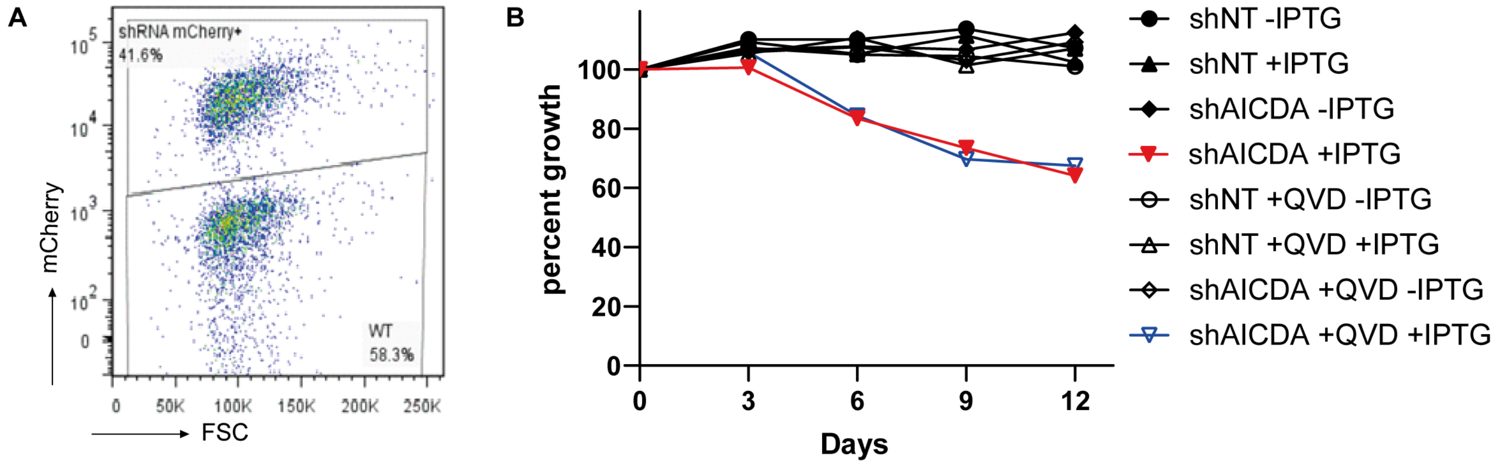


**Supplementary Figure 3. AID inhibition decreases proliferative fitness of TMD8 cells in a competition setting.**
**A**, Flow cytometry plots illustrating two populations in the competition assay, in which mCherry-positive shRNA-expressing clones were co-cultured with mCherry-negative wild-type (WT) cells. Forward scatter (FSC) is shown on the x-axis and mCherry fluorescence on the y-axis. **B**, Growth curves of competition assay in which shAICDA- or shNT-expressing cells were mixed with WT cells in triplicate in the presence or absence of IPTG (to induce shRNA expression) and QVD (to inhibit apoptosis). Cells were seeded and maintained at 0.3 × 10^6 cells/ml with passaging every 3 days. At each passage, cell populations were quantified by flow cytometry and normalized to the initial seeding ratio with WT cells.

Supplementary Figure 4

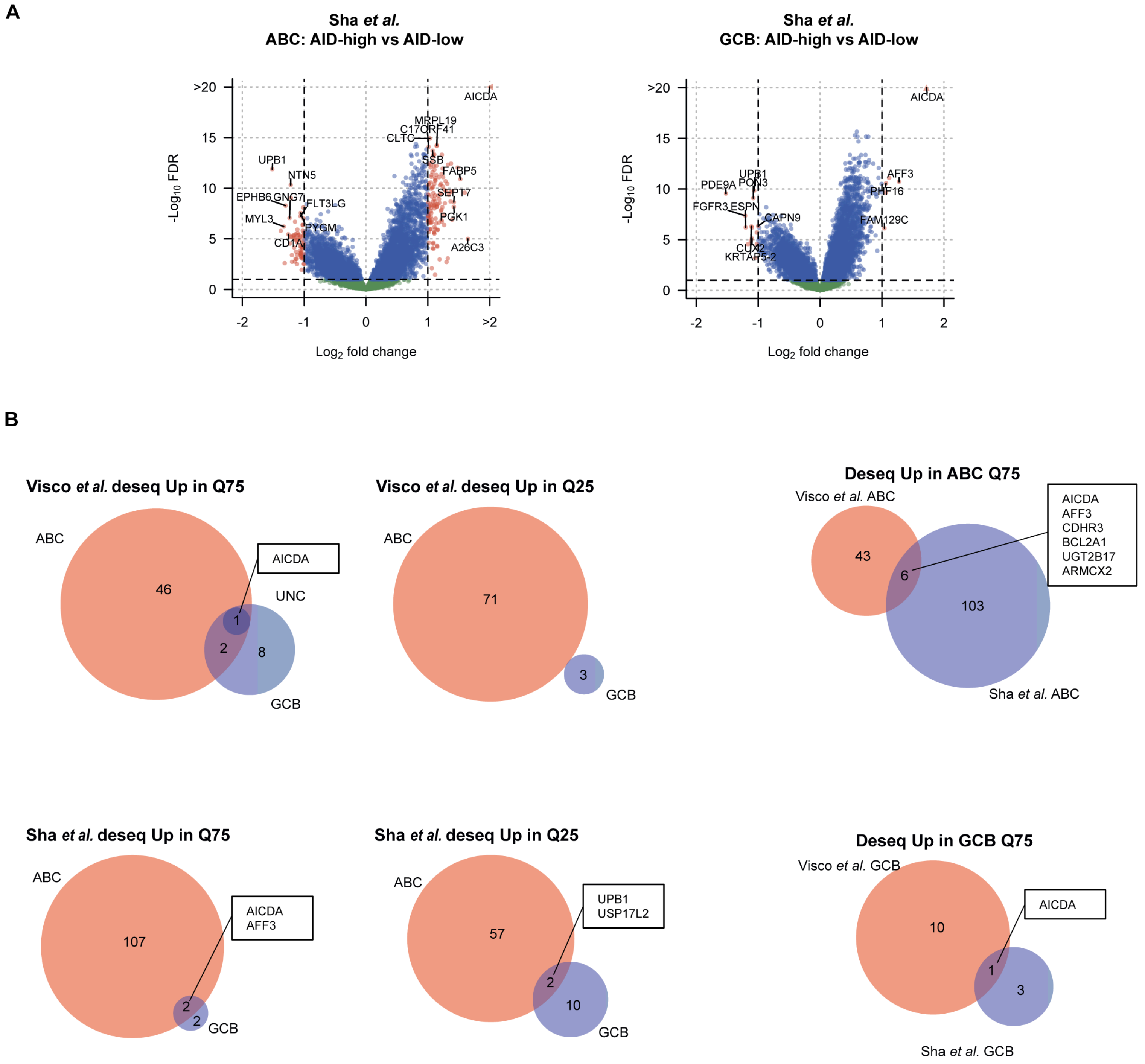


**Supplementary Figure 4. Differential gene expression in DLBCL patients based on AID expression.**
**A**, Volcano plots showing DESeq2 differential gene expression analysis on ABC-type (left panel) and GCB-type (right panel) DLBCL patient samples from Sha *et al.* comparing the upper and lower quartiles of AID expression. Significance was defined as FDR<0.1 and log(FC) <-1 and >1. **B**, Venn diagrams showing differentially expressed genes in the upper and lower quartiles of AID expression in the distinct DLBCL subgroups from the Visco *et al.* dataset (top left two panels) and the Sha *et al.* dataset (bottom left two panels). The right panels show overlap of differentially expressed genes within each DLBCL subtype across patient groups in the upper quartile of AID expression: ABC-type (top right) and GCB-type (bottom right).

Supplementary Figure 5

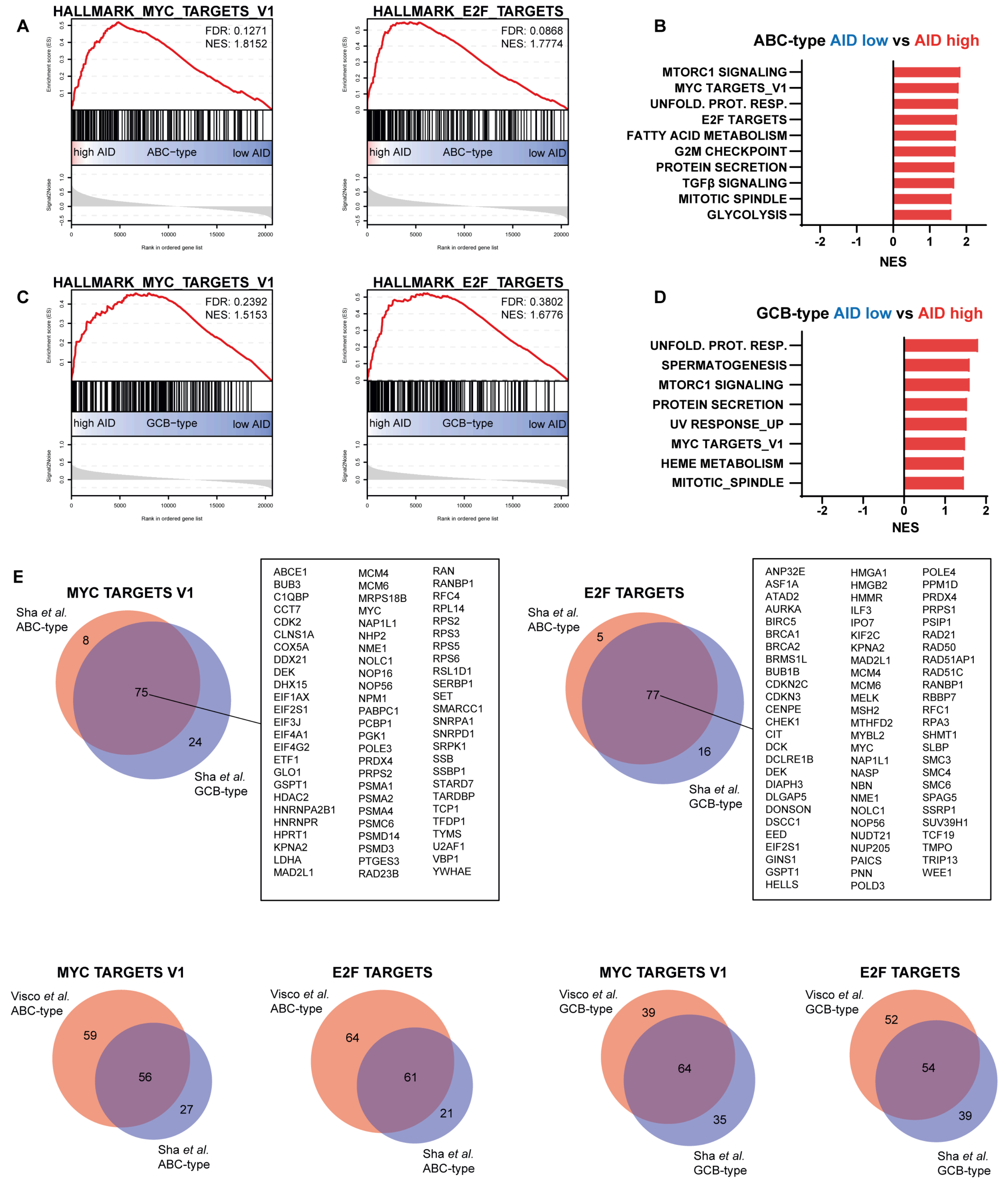


**Supplementary Figure 5. High AID expression is associated with enhanced MYC and E2F pathway activity in DLBCL.** **A-D**, GSEA enrichment plots of MYC target genes (left panels) and E2F target genes (right panels) in ABC-type (**A**) and GCB-type (**C**) DLBCL patient samples from Sha *et al.*. Normalized enrichment scores (NES) and false discovery rates (FDR) are indicated in each plot. Enriched pathways are summarized in bar graphs for ABC-type (**B**) and GCB-type (**D**) DLBCL. In ABC-type (**B**), pathways with FDR < 0.2 are shown. In GCB-type (**D**), no pathways achieved FDR < 0.2, and thus pathways with FDR < 0.25 are displayed. **E**, Venn diagrams showing the overlap of MYC target genes (top left) and E2F target genes (top right) between ABC- and GCB-type DLBCL patients from Sha *et al.*. The lower panels depict overlap of MYC (first and third panels) and E2F (second and fourth panels) target genes across patient groups from Visco *et al.* and Sha *et al.*, stratified by DLBCL subtype: ABC-type

Supplementary Figure 6

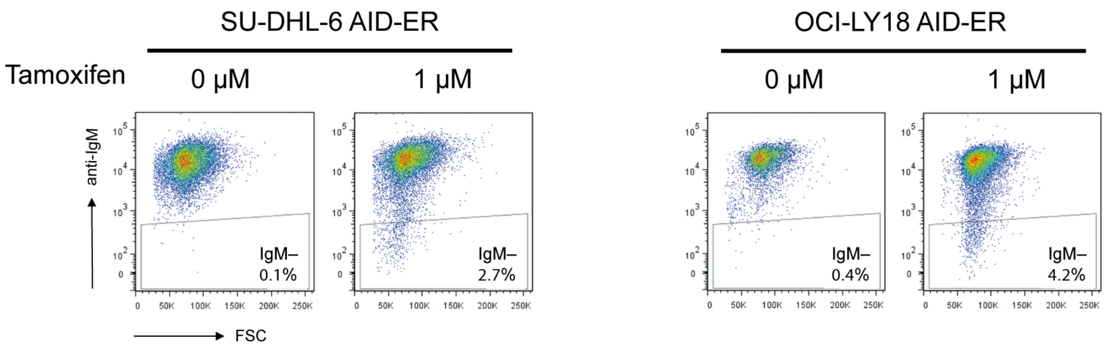


**Supplementary 6: AID-ERα activation induced with tamoxifen is shown by surface IgM loss in GCB-type DLBCL cell lines SU-DHL-6 and OCI-LY18.**

Flow cytometry plots showing surface IgM expression in GCB-type DLBCL cell lines SU-DHL-6 (left panel) and OCI-LY18 (right panel) AID-ERα overexpressing cell lines cultured for 12 days in the presence or absence of tamoxifen. Forward scatter (FSC) is shown on the x-axis and surface IgM expression on the y-axis.

Supplementary Figure 7

**A**

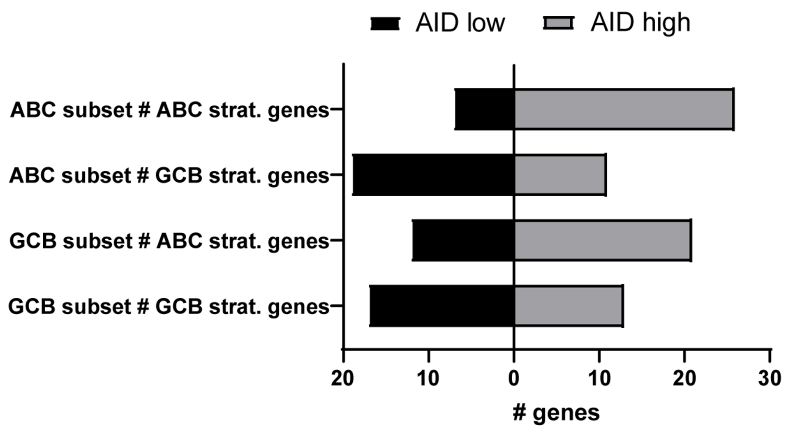


**B**

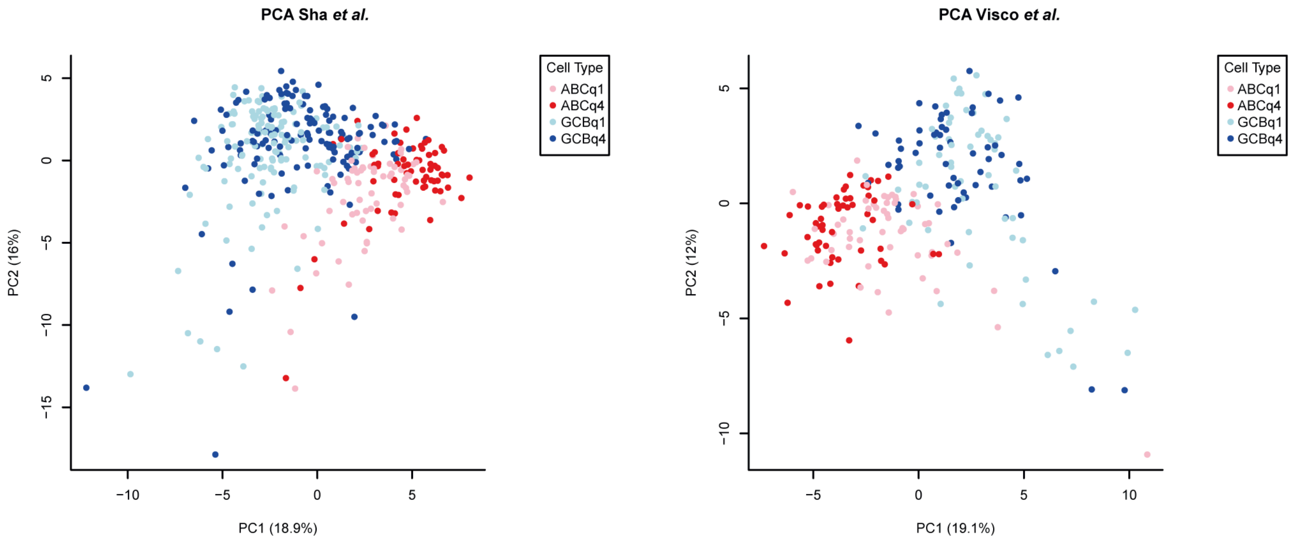


**Supplementary 7. AID expression promotes an ABC-like gene expression profile in ABC- and GCB-type DLBCL patients. A**, Bar graphs showing the number of ABC and GCB stratification genes with higher expression in the upper quartile versus the lower quartile of AID expression (grey) or higher expressed in the lower quartile than the upper quartile of AID expression (black), stratified by ABC-type and GCB-type DLBCL patients from Visco *et al.*. **B**, Principal component analysis (PCA) of stratification genes was performed using log2 z-scores in four subgroups from the Sha *et al.* dataset (left panel) and the Visco *et al.* dataset (right panel): GCB AID low (GCBq1), GCB AID high (GCBq4), ABC AID low (ABCq1), and ABC AID high (ABCq4).
